# The Disruption and Normalization of Cortical Language Network Dynamics After Post-Stroke White Matter Disconnection

**DOI:** 10.1101/2025.08.25.672078

**Authors:** Tao Zhong, Qingwen Chen, Xiaolin Guo, Junjie Yang, Xiaowei Gao, Zhe Hu, Jian Liu, Junjing Li, Jiaxuan Liu, Yaling Wang, Zhiheng Qu, Wanchun Li, Zhongqi Li, Wanjing Li, Yien Huang, Jiali Chen, Hao Wen, Binke Yuan, Han Gao

**Author notes:** **Correspondence:** Corresponding Author Han Gao, M.D, Department of Neurosurgery, The Affiliated Qingyuan Hospital (Qingyuan People’s Hospital), Guangzhou Medical University, Qingyuan, China, Address: No.35, Yinquan North Road, Qingcheng District, Qingyuan Guangdong, China, Postcode: 511518, Binke Yuan, Ph.D, Room 206, Institute for Brain Research and Rehabilitation, South China Normal University, Guangzhou, China, Postcode: 510631. These authors contributed equally to this work (Tao Zhong, Qingwen Chen, Xiaolin Guo).

## Abstract

The human language network undergoes reorganization across different spatiotemporal scales, limited by its fixed structural neurobiological foundations. Although stroke often damages white matter, its effect on temporary language network reorganization is not fully understood. This study examined longitudinal behavioral and resting-state fMRI data from post-stroke patients with exclusive subcortical lesions at three time points: two weeks (n = 38), three months (n = 29), and one year (n = 27). Patients showed mild to moderate language impairments during the acute phase, which improved within three months. The extent of disconnection in several left-hemispheric white matter tracts was negatively associated with language deficits. Healthy controls (HC, n = 25) exhibited domain-segregation dynamics in the cortical language network, limited by the underlying white matter pathways. In patients, severe, state- and track-specific network disruptions were observed in the acute phase, characterized by hypo- and hyper-connectivity and abnormal topological features. As language recovery progressed, connectivity patterns began to return to normal, resembling those of HC, and the domain-segregation dynamics reappeared. These results deepen our understanding of structure-function coupling and indicate significant post-lesional plasticity within the cortical language network.

## Introduction

Language processing depends on a network of distributed gray matter regions and the white matter pathways connecting them (Duffau, Moritz-Gasser, & Mandonnet, 2014). Brain areas involved in language processing extend laterally across the frontal, temporal, and parietal cortex (Fedorenko & Thompson-Schill, 2014; Hickok, 2022; Price, 2012). These regions are connected by white matter pathways including the arcuate fasciculus (AF), inferior longitudinal fasciculus (ILF), inferior occipital-frontal fasciculus (IFOF), middle longitudinal fasciculus (MdLF), uncinate fasciculus (UF), and frontal aslant tract (FAT) (Duffau et al., 2014; Saur et al., 2008; Yeh, 2022). Within this relatively stable structural framework, the cortical language network dynamically reorganizes across multiple spatial and temporal scales. Spatially, the language network is arranged into hierarchical, segregated large-scale subnetworks, with a core dorsal (phonological) stream and ventral (semantic) stream (Hickok, 2022; Lu et al., 2021; Price, 2012; Vigneau et al., 2006; B. Yuan, Xie, Wang, et al., 2023). Temporally, these specialized regions are involved in expressive and receptive language tasks in a hierarchical and sequential manner, with activity captured over time scales ranging from milliseconds to minutes (Abbasi, Steingraber, Chalas, Kluger, & Gross, 2023; Chai, Mattar, Blank, Fedorenko, & Bassett, 2016; He et al., 2018; Liljestrom, Kujala, Stevenson, & Salmelin, 2015; Liu et al., 2020; Youssofzadeh & Babajani-Feremi, 2019).

Extensive research shows that focal brain damage, such as stroke, glioma, and traumatic brain injury, can cause language impairments if language-related cortical regions or white matter tracts are impacted (Corbetta et al., 2015; Marini et al., 2011; B. Yuan, Xie, Gong, et al., 2023; Zhang et al., 2021). Neuronal death leads to local brain function loss, while white matter disconnection disrupts information exchange between connected regions. At the same time, the brain functions as an integrated system, with mutual influence between gray and white matter: damage to one affects the other’s functionality (Honey et al., 2009; Langen et al., 2017; Messe, 2020; Nozais et al., 2023; van den Heuvel, Mandl, Kahn, & Hulshoff Pol, 2009; Wang, Dai, Gong, Zhou, & He, 2015). However, because of the pathological features of these diseases, lesions often involve both white and gray matter, making it difficult to determine whether a patient’s language impairment results from neuronal damage or white matter tract disruption (Favaretto et al., 2022; Griffis, Metcalf, Corbetta, & Shulman, 2019; Nozais et al., 2023; Parsons, Hughes, Poudel, D., & Caeyenberghs, 2020; Salvalaggio, De Filippo De Grazia, Zorzi, Thiebaut de Schotten, & Corbetta, 2020).

In this study, we examined patients after pure subcortical stroke to understand how white matter disconnection influences the transient reconfiguration of the cortical language network. At two weeks, three months, and one year post-stroke, we collected resting-state functional MRI (rs-fMRI) data and language performance scores. For each patient, we measured the degree of disconnection across all white matter tracts based on the damage area in the acute phase. Using a cortical language network atlas containing 68 regions of interest (ROIs), we applied the dynamic conditional correlation (DCC) method to create each participant’s framewise dynamic functional connectivity matrices. We then conducted a cluster analysis to identify recurring temporal states. Finally, we evaluated how white matter disconnection impacted the dynamic reorganization of the cortical language network and the recovery process.

Unlike previous studies on data-driven dynamic brain networks, this research used the dynamic meta-networking framework of language (B. Yuan, Xie, Wang, et al., 2023) to examine the dynamic features of the cortical language network after subcortical stroke. Specifically, the dynamic meta-networking framework of language is a theoretical model of language network activity during rest, consisting of four recurring states with distinct connectivity patterns, hub locations, structural bases, and cognitive significance. In state 1, regions in the bilateral superior temporal gyrus (STG) and ventral parts of precentral gyrus (PrG) and postcentral gyrus (PoG) served as densely connected hubs. In state 2, hubs are mainly found in the prefrontal and posterior temporal cortex. In state 3, hubs are primarily located in the temporal cortex. The fourth state lacks hubs and is characterized by weak functional connectivity. The white matter connections underlying these hubs in the first three states influenced their distribution. Spatially, the hub patterns in these states closely correspond to neural processes involved in speech perception, lexical-phonological processing, speech production, and semantic processing. These four states together form a dynamic “meta-networking” framework for language. Meta-networking refers to a network of interconnected networks, a central concept in the “meta-networking” theory of brain functions (Herbet & Duffau, 2020). This theory suggests that complex cognition and behaviors (such as language) originate from the spatiotemporal integration of distributed but specialized sub-networks. The dynamic “meta-networking” framework captures the domain-specific nature of language processing, aligns with the neurobiology outlined by the dual-stream model of speech and language, and has demonstrated relevance in cognitive and clinical contexts (B. Yuan, Xie, Gong, et al., 2023).

Based on this framework, we can form the following hypotheses. First, damage to cortical language areas and the underlying white matter tracts can disrupt this optimized system. Second, different damage locations, whether in white matter or cortical areas, may cause pathway-or state-specific impairments. Finally, pathway- or state-specific damage might be linked to particular levels of structured language deficits. To avoid the confounding effects of cortical damage, this study only included stroke patients with subcortical white matter injury. This method helps examine how white matter disconnection influences the dynamic reorganization of cortical language networks.

## Materials and Methods

The stroke dataset we used is part of the Washington Stroke Cohort, which is publicly available at https://cnda.wustl.edu/data/projects/CCIR_00299. The dataset includes longitudinal structural imaging, resting-state fMRI, and neuropsychological testing scores at two weeks, three months, and one year after stroke. Written informed consent was obtained from all participants following the Declaration of Helsinki and procedures established by the Washington University in Saint Louis Institutional Review Board.

### Subjects

The entire dataset included 132 patients with their first symptomatic stroke, and their data had been previously published using different approaches from those used in this study (Corbetta et al., 2015). We selected patients with subcortical stroke based on lesion masks created by neurosurgeons C.Q. and Z.T. Ultimately, 38 patients at two weeks, 29 at three months, and 27 at one year post-stroke met the inclusion criteria. Additionally, 25 healthy controls (HCs), matched for age, sex, and education, were recruited.

### MRI imaging

MRI imaging data was acquired with a Siemens 3T Tim-Trio scanner at the School of Medicine at Washington University in St. Louis. Structural scans included: (1) a sagittal MPRAGE T1-weighted image with TR=1950 ms, TE=2.26 ms, flip angle=9°, voxel size=1.0 × 1.0 × 1.0 mm, and slice thickness=1.00 mm; (2) a transverse turbo spin-echo T2-weighted image with TR=2500 ms, TE=435 ms, voxel size=1.0 × 1.0 × 1.0 mm, and slice thickness=1.00 mm; and (3) a sagittal FLAIR, fluid-attenuated inversion recovery, with TR=7500 ms, TE=326 ms, voxel size=1.5 × 1.5 × 1.5 mm, and slice thickness=1.50 mm. Resting-state functional scans were obtained using a gradient echo EPI sequence (TR=2000 ms, TE=27 ms, 32 contiguous 4-mm slices, 4 × 4 mm in-plane resolution), during which participants were instructed to fixate on a small white cross centered on a black background screen in a low luminance environment. Six to eight resting-state (RS) fMRI runs, each consisting of 128 volumes, were collected.

### Lesion drawing and disconnection severities of white matter tracts

For each patient, lesions were manually delineated on T1-weighted images obtained two weeks after stroke, with T2-weighted images and FLAIR images used as visual references. The native stroke mask was spatially normalized to standard Montreal Neurological Institute (MNI) space by applying the deformation field estimated through tissue segmentation of T1-weighted images. The percentage of white matter tract disconnection was estimated using the Lesion Quantification Toolkit (Griffis, Metcalf, Corbetta, & Shulman, 2021). This toolkit estimates tract disconnection severity in the HCP-1065 population-averaged streamline tractography atlas (Yeh, 2022), which was generated from the tractograms of 1065 subjects in MNI space and includes 91 canonical macroscale white matter tracts (Supplementary Table 1). To assess disconnection severity for each tract, the lesion is first embedded into the HCP-1065 streamline tractography atlas as a region of interest (ROI), then intersected with the streamlines of the 91 tracts. The number of disconnected streamlines for each tract is converted into a percentage of the total streamlines assigned to that tract, with higher values indicating more severe white matter disconnection.

### Rs-fMRI preprocessing

The resting-state fMRI data were preprocessed with the following steps: 1) remove the first 10 volumes; 2) apply slice timing correction; 3) perform head motion correction (< 3 mm or 3 degrees); 4) coregister images; 5) segment T1 images and normalize them into the MNI space using DARTEL; 6) normalize functional images using the deformation field from T1 images; 7) smooth data with a Gaussian kernel (full-width-at-half-maximum = 6 mm); 8) apply linear detrending; 9) regress out nuisance signals (24 parameters, including *x, y, z* translations and rotations (6 parameters), plus their derivatives (6 parameters) and quadratic terms of 12 parameters); 10) perform temporal band-pass filtering (0.01–0.1 Hz).

### The cortical language network definition

To investigate the cortical language network dynamics, we defined a putative cortical language network according to the Human Brainnetome Atlas (Fan et al., 2016). We extracted all parcels that exhibited significant activation in language-related behavioral domains (i.e., language or speech) or paradigm classes (e.g., semantic, word generation, reading, or comprehension). Thirty-three cortical parcels in the left hemisphere and 15 in the right hemisphere were selected. Their extents are highly similar to those identified by other researchers (Lipkin et al., 2022; Vigneau et al., 2006). Considering bilateral language processing and the recruitment of right hemisphere language regions for recovery, 19 homologs in the right hemisphere and one parcel of homologs in the left hemisphere were also selected. Altogether, a symmetric language network including 68 cortical ROIs (34 ROIs in each hemisphere) was defined (Binke Yuan et al., 2022), including the superior, middle, and inferior frontal gyrus (SFG, MFG, and IFG, respectively), the ventral parts of PrG and PoG, STG, middle and inferior temporal gyrus (MTG, ITG), the fusiform gyrus (FuG), the Para hippocampus gyrus (PhG) and the posterior superior temporal sulcus (pSTS).

### Frame-wise construction of time-varying networks using DCC

To identify the recurring temporal states of two types of networks, the dynamic conditional correlation (DCC) approach was used to build the framewise, time-varying language network (Lindquist, Xu, Nebel, & Caffo, 2014; Yang et al., 2025).

DCC is a variant of the multivariate GARCH (generalized autoregressive conditional heteroscedasticity) model (Lebo & Box‐Steffensmeier, 2008), which is especially effective for estimating both time-varying variances and correlations. GARCH models represent the conditional variance of a single time series at time t as a linear combination of past values of the conditional variance and the squared process itself. All the parameters of DCC are estimated using quasi-maximum likelihood methods and do not require any ad hoc parameter settings.

The DCC algorithm consists of two steps. To illustrate, let us assume that there is a pair of time series from two ROIs, *x*_*t*_ and *y*_*t*_. In the first step, standardized residuals of each time series are estimated using a univariate GARCH (1,1) process. In the second step, an exponentially weighted moving average (EWMA) window is applied to the standardized residuals to compute a non-normalized version of the time-varying correlation matrix between *x*_*t*_ and *y*_*t*_. The mathematical expressions of the GARCH (1,1) model, DCC model, and EWMA, and the estimations of the model parameters were provided by Lindquist et al. (2014).

K-means clustering was used to decompose the dFC matrices into several recurring connectivity states. The optimal number of clusters, k, was determined using the elbow criterion, which considers the ratio between the within-cluster distance and the between-cluster distances (Allen et al., 2014; Yang et al., 2025). The L1 distance function, also known as ‘Manhattan distance,’ was employed to measure the distance from points to centroids. Each time window was ultimately assigned to one of these connectivity states.

### Topological properties of dFC states

A state-specific subject connectivity matrix was created by taking the median of all connectivity matrices associated with the same state label (Damaraju et al., 2014). Before calculating, a correlation threshold (*r* > 0.2) was applied to remove weak connections likely caused by noise. Because of the controversial nature of negative connections (Murphy & Fox, 2017), only positive connections were included in this work. In this study, we used a weighted network instead of a binarized one to preserve all connectivity information.

#### Global topological metrics

three network topological properties were computed for each state-specific matrix: 1) Total connectivity strength. The FC strength of a network was calculated by summing the functional connectivity strengths of all suprathreshold connections into one value; 2) Global network efficiency (gE). The gE indicates the ability for parallel information transfer and functional integration. It is defined as the average of the inverses of all weighted shortest path lengths (the minimal number of edges that one node must traverse to reach another) in the thresholded matrix (Rubinov & Sporns, 2010); 3) Local network efficiency (lE). The lE reflects the relative functional segregation and is defined as the average global efficiency of each node’s neighborhood sub-graph.

#### Nodal topological metrics

We calculated each node’s nodal strength, gE, and lE. We also calculated betweenness centrality, an index indicating whether a specific node lies on the shortest paths between all pairs of nodes in the network.

### Functional significance of state-dependent hub distributions

To evaluate the functional importance of hubs in each state, we conducted term-based meta-analyses using the NeuroQuery platform (https://neuroquery.org/) (Dockes et al., 2020). NeuroQuery predicts the spatial distribution of activity from 418,772 activation locations associated with 7647 terms in the full texts of 13,459 neuroimaging articles. To assess functional specificity, we analyzed five terms— speech perception, inner speech, phonological processing, speech production, and semantic processing. The functional relevance of hubs was measured by calculating dice coefficients between binary images of hub nodes and the meta-analysis results using the formula: dice = 2 ^*^ (hub ∩ meta) / (hub + meta). Due to the left-lateralized nature of the meta-analysis activations, dice coefficients were calculated specifically for the left hemisphere.

### Statistical analysis

Age data were analyzed using Two-sample t-tests. Sex data were using Pearson’s chi-square test.

For edge strength along with nodal and global topological properties, two-sample t-tests were conducted. The results for edge strength were corrected using network-based statistics (NBS) (Zalesky, Fornito, & Bullmore, 2010) with an edge P-value of 0.05 and a component P-value of 0.05. The results for nodal and global topological properties were corrected using False Discovery Rate (FDR) with a corrected P-value of 0.05. The associations between edge, nodal, and global topological properties and patients’ language scores were assessed by calculating the partial Pearson correlation coefficient after controlling for age, gender, lesion type (including Ischemic and Hemorrhagic), race, and education. The relationship between edge strength and language scores was also corrected using NBS with a component P-value of 0.05.

## Results

### Lesion anatomy, white matter disconnections, and language deficits

Most patients had circumscribed stroke lesions in the cortical white matter, while several patients had stroke lesions in the white matter of the brain stem and cerebellum. The highest lesion overlaps were in the bilateral superior corona radiata (Figure 1).

**Figure 1.**
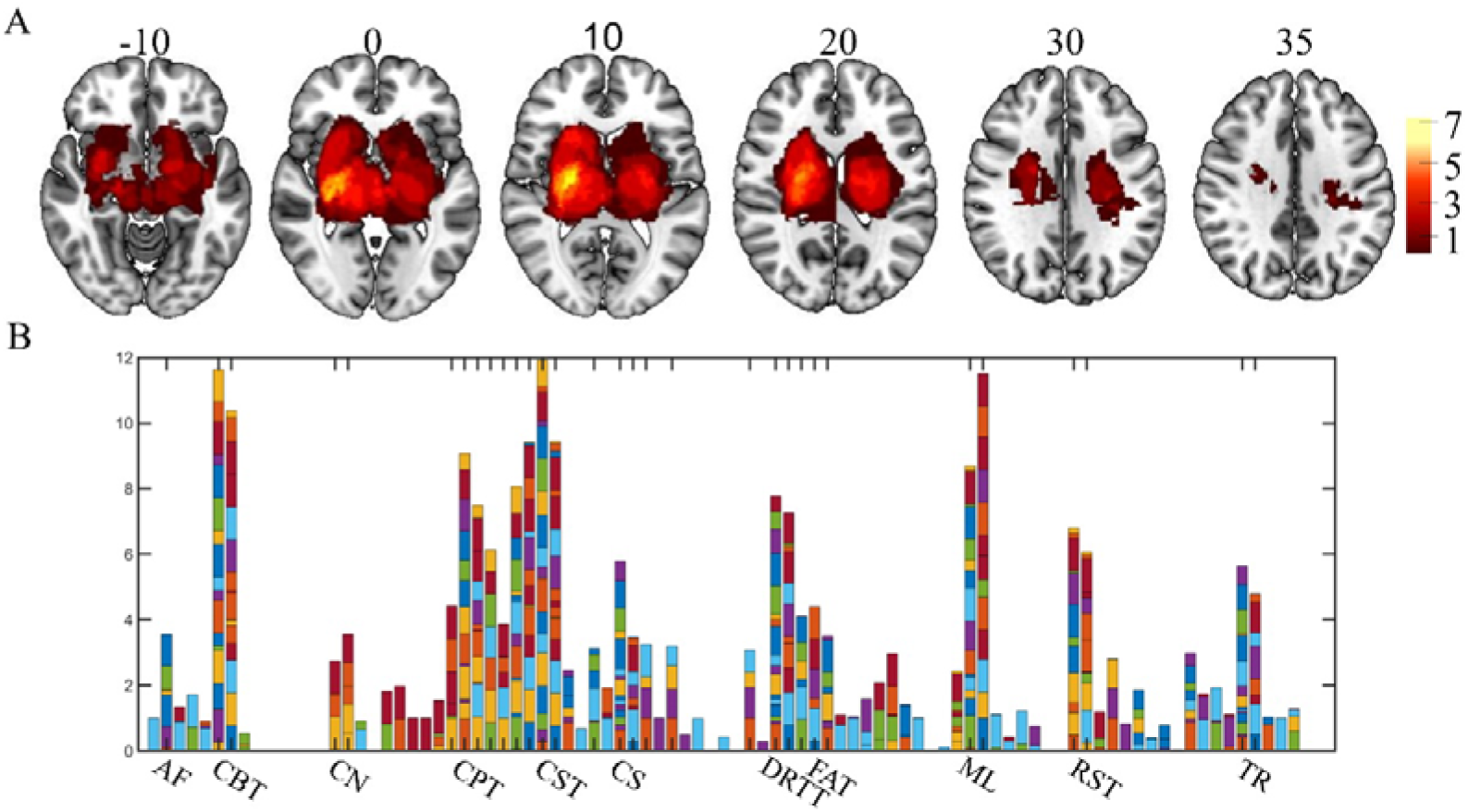
Lesion anatomy and white matter disconnection. A: Lesion overlap map showing 38 patients within two weeks after stroke. The color bar indicates the number of patients with lesions in each specific voxel. B: The number of white matter disconnections for each patient. A group-averaged tractography atlas with 91 white matter tracks, specifically the HCP-1065 (Yeh, 2022), was used to quantify white matter disconnections. AF, Arcuate Fasciculus; CBT, Corticobulbar Tract; CN, Cranial Nerves; CPT, Corticopontine Tract; CST, Corticospinal Tract; CS, Corticostriatal Tract; DRTT, Dentatorubrothalamic Tract; FAT, Frontal Aslant Tract; ML, Medial Lemniscus; RST, Reticulospinal Tract; TR, Thalamic Radiation.

Sets of white matter tracks were damaged, including the AF, corticobulbar tract (CBT), cranial nerves (CN), corticopontine tract (CPT), corticospinal tract (CST), corticostriatal tract (CS), dentatorubrothalamic tract (DRTT), frontal aslant tract (FAT), medial lemniscus (ML), reticulospinal tract (RST), and thalamic radiation (TR). Patients exhibited significantly higher blood pressure than HCs (Table 1). Language deficits appeared two weeks after a stroke, mainly due to severe impairments in complex ideational tasks, oral sentence reading, and stem completion (Figure 2 and Table 2). These language deficits mostly recovered within three months post-stroke.

**Table 1.** Demographic and clinical information of stroke patients and healthy controls

| Variables | Controls(n=25) | Stroke |  |  | *P* value |
| --- | --- | --- | --- | --- | --- |
|  |  | 2W(n=38) | 3M(n=29) | 1Y(n=27) |  |
| Age | 55.84±10.82 | 54.74±9.59 | 55.76±7.30 | 57.15±5.73 | 0.608 |
| Gender(female/male) | 14/11 | 16/22 | 12/17 | 10/17 | 0.280 |
| Lesion_type |  |  |  |  | - |
| Ischemic | - | 34(89%) | 25(86%) | 25(93%) |  |
| Hemorrhagic | - | 4(11%) | 4(14%) | 2(7%) |  |
| Race |  |  |  |  | 0.254 |
| White | 10(40%) | 10(26%) | 6(21%) | 5(19%) |  |
| Black or African American | 15(60%) | 28(74%) | 23(79%) | 22(81%) |  |
| Years of education | 13.24 ±2.22 | 13.24 ±2.52 | 13.24 ±2.53 | 13.78 ±2.71 | 0.886 |
| Smoke | 10(40%) | 24(63%) | 19(66%) | 17(63%) | 0.071 |
| High blood pressure | 0(0%) | 27(71%) | 21(72%) | 22(81%) | <0.001 |
| Diabetes | 3(12%) | 12(32%) | 10(34%) | 10(37%) | 0.074 |

Gender, Lesion type, Race, Stroke, High blood pressure, and diabetes are measured using a Chi-squared test, and age and education are measured using two-sample t-tests.

**Table 2.** Results of linear mixed-effects model for language tests

|  | P-value |  |  |
| --- | --- | --- | --- |
|  | Group | Post-stroke Time | Group&Time |
| Basic word discrimination | 0.167 | 0.256 | 0.861 |
| Commands | 0.149 | 0.144 | 0.553 |
| Complex ideational material | **<0.001** | **<0.001** | 0.212 |
| Boston naming test short form | 0.251 | 0.113 | 0.271 |
| Oral reading of sentences | **0.019** | 0.038 | 0.612 |
| Comprehension of oral reading of sentences | 0.107 | 0.455 | 0.518 |
| Non-word reading | 0.074 | 0.174 | 0.932 |
| Stem completion | **0.005** | **0.046** | 0.879 |
| Animal naming | 0.393 | 0.534 | 0.078 |
| PCA Score | **0.027** | 0.083 | 0.890 |

**Figure 2.**
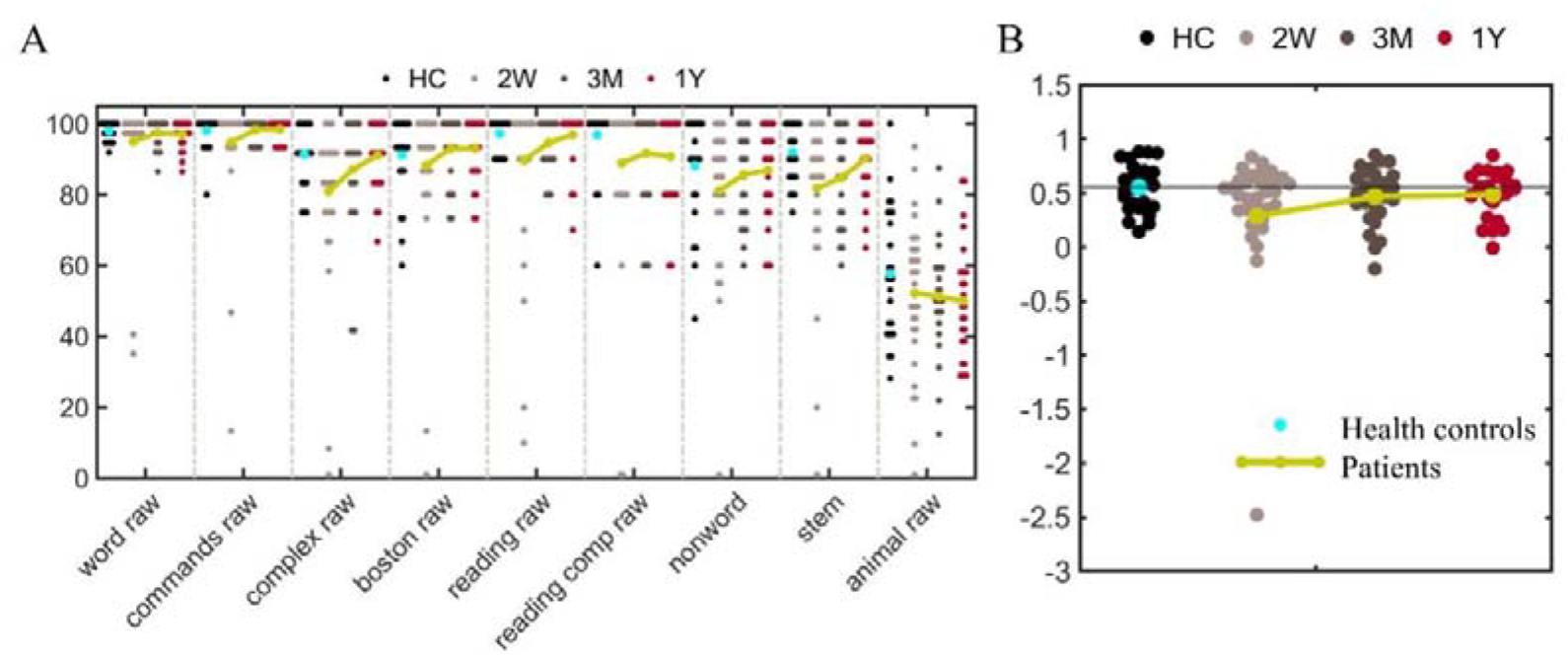
Language deficits during the acute phase and recovery.

The levels of white matter disconnection were significant (*P* < 0.05, FDR correction) and negatively associated with language deficits during the acute phase (Table 3). The strongest association is seen in the anterior and middle regions of the corpus callosum. The other connected fiber tracts are all situated in the left hemisphere, including language-related tracts such as the IFOF, MdLF, FAT, and AF, as well as non-language tracts like the superior longitudinal fasciculus, fornix, extreme capsule, and others.

**Table 3.**
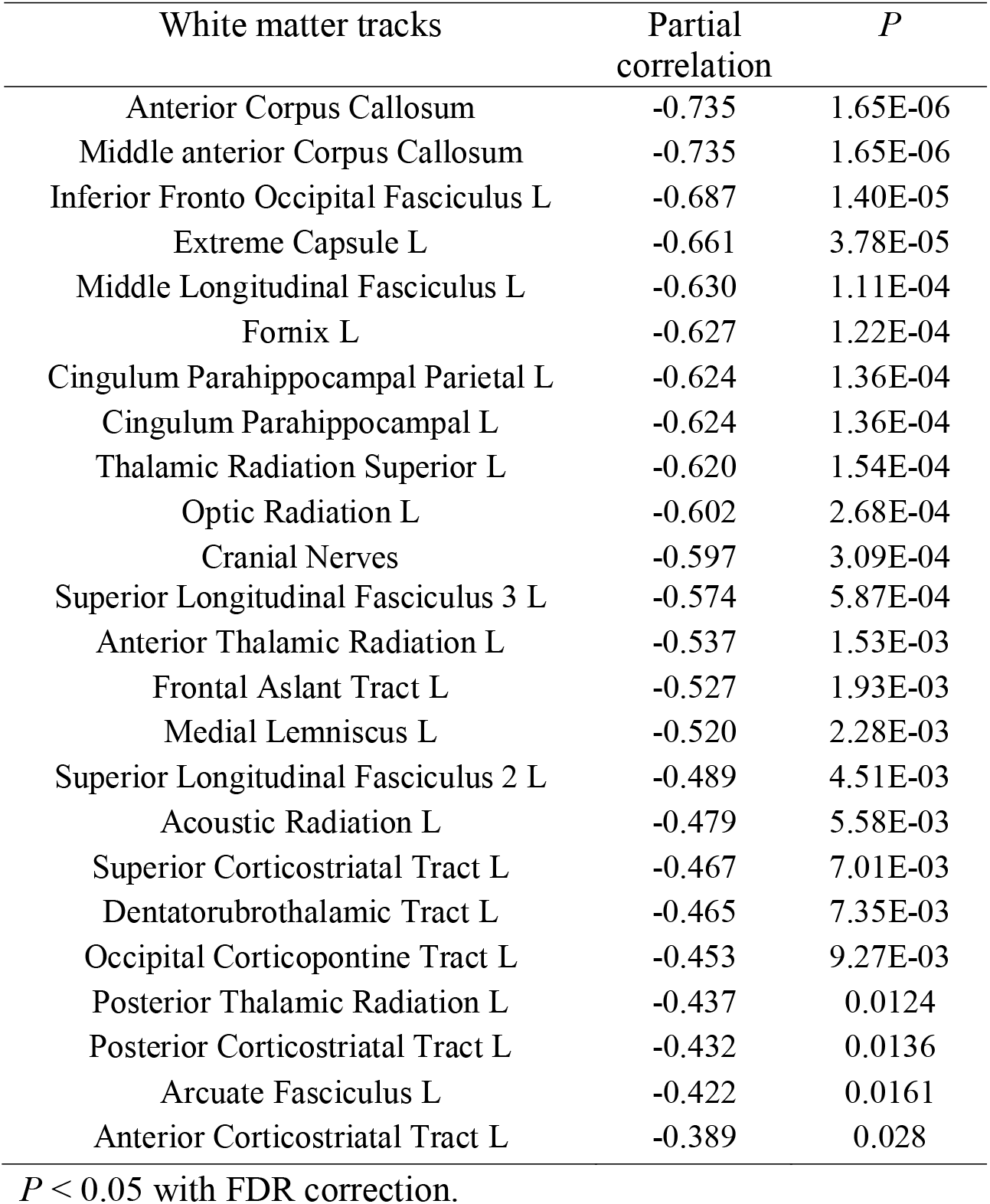
Associations between the white matter disconnected severities and language deficits.

### The disruptions in cortical language network dynamics and their normalization during language recovery

Four recurring temporal states with distinct connectivity patterns and hub distribution were identified in both healthy controls (HCs) and patients (Figures 3 and 4). The nodal strength distributions of all states were best described by the exponentially truncated power-law form (Figure 3 and Supplementary Figure 1), indicating the presence of a small number of highly connected hub nodes. In each group, the top 20 nodes with the highest nodal strengths were defined as hubs (Figures 3 and 6). We arranged the states based on their hub distributions.

**Figure 3.**
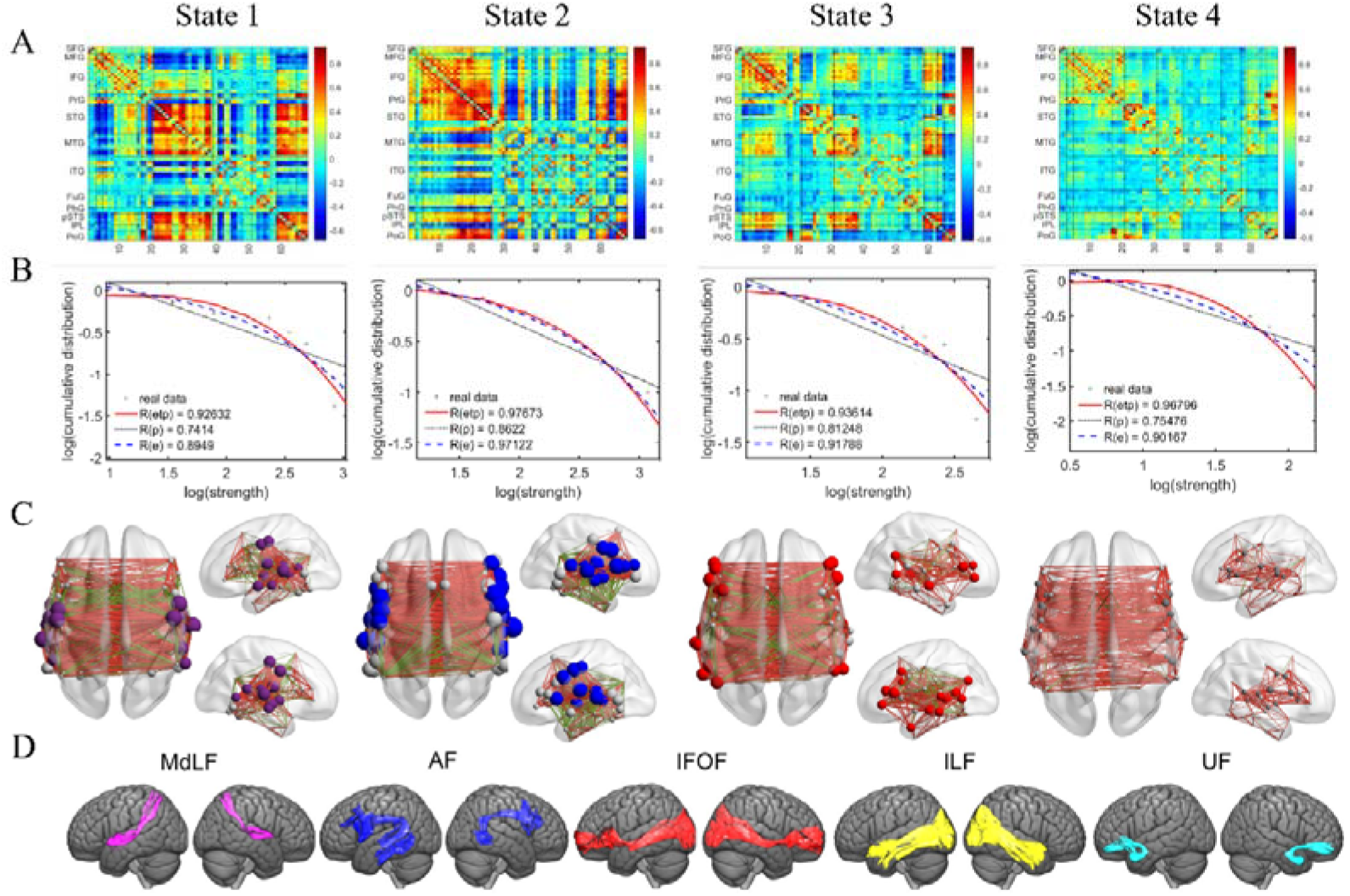
The domain-segregation cortical language network dynamics in HCs. A: Four recurring temporal states with distinctive functional connectivity were identified. B: Nodal strength distribution. The log-log plots show the cumulative nodal strength distributions in patients. The plus sign (black) represents observed data, and the solid red line is the fit of the exponentially truncated. power-law, *P* (*x*) ∼ *x*^α − 1^exp (*x*/*x*_*c*_), the dashed line (blue) is an exponential, *P* (*x*) ∼ exp (*x*/*x*_*c*_), and the dotted line (black) is a power-law, *P* (*x*) ∼ *x*^α − 1^.*R*^2^ was calculated to assess the goodness-of-fit. A larger value indicates a better fit: *Retp, R*^*2*^ for the exponentially truncated power-law; *Re, R*^*2*^ for the exponential; Rp, *R*^*2*^ for the power-law fit. The exponentially truncated power-law is the best fit for all four states, suggesting long-tailed broad-scale topologies and that a large proportion of network connectivity is concentrated on a subset of nodes (i.e., hubs). C: The first 20 nodes with the highest nodal strength in each state are colored and defined as hubs. D: Structural neurobiology of hubs. Language-related white matter bundle terminations provided by Yeh (2022). MdLF: middle longitudinal fascicle, AF: the arcuate fascicle, ILF: inferior longitudinal fascicle, IFOF: inferior fronto-occipital fasciculus, and UF: uncinate fasciculus.

**Figure 4.**
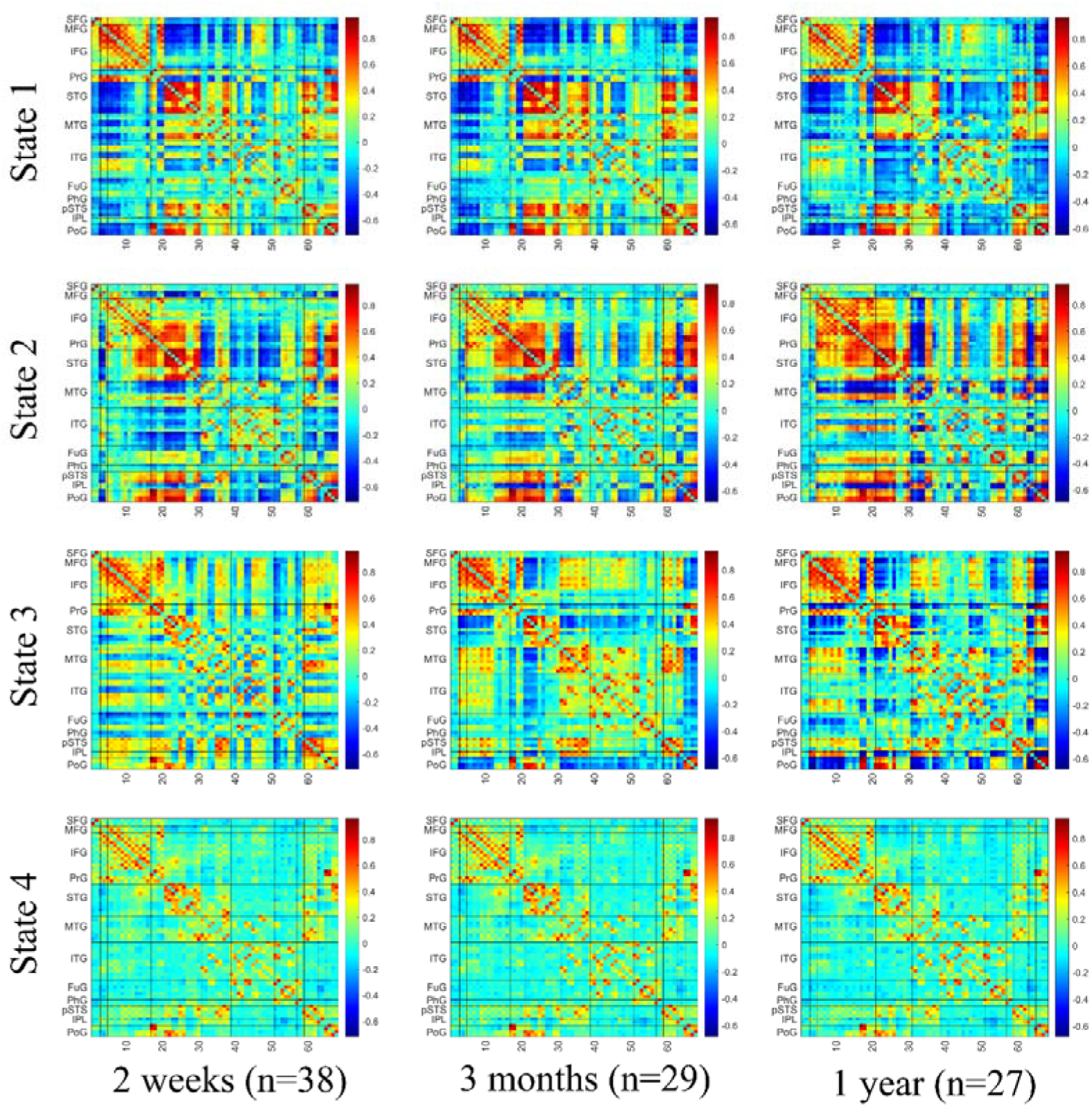
The recurrent states over time in patients.

In state 1, hubs were mainly located in STG. In state 2, hubs were primarily found in the prefrontal and posterior temporal cortex. In state 3, hubs were mainly situated in the temporal cortex, posterior parietal cortex, and IFG. By analyzing the spatial similarity between term-based activation and hub distribution, we observed that hubs in the first three states highly overlapped with activation related to speech perception, inner speech, phonological processing, and speech production, and partially overlapped with semantic processing (Supplementary Figure 2). The functional significance of the hub distributions in these three states revealed a domain-segregation pattern in cortical language network dynamics.

In state 1, the tracks connecting hubs are MdLF and AF. In state 2, the white matter tracks beneath the hubs are the frontal parts of AF and the temporal parts of IFOF. In state 3, the white matter tracks beneath the hubs are ILF, IFOF, and the temporal parts of AF.

The dynamics of the domain-segregation cortical language network were observed in healthy controls (HCs) and patients at three months and one year after stroke. Two weeks post-stroke, reduced spatial correlations and weak dynamics were seen in states 2 and 3 (Figure 5). Hubs in these states reappeared during language recovery. Statistical analysis indicated that the functional connectivity in the first three states was heavily disrupted two weeks after stroke. Both decreased and increased connectivity were noted (Figure 7). Interhemispheric connections mainly drive these hypo- and hyper-connectivity patterns. Partial correlation analysis found significant positive and negative relationships between hypo- and hyper-connectivity and language performance, respectively, suggesting that connectional diaschisis has dual effects (Figure 8).

**Figure 5.**
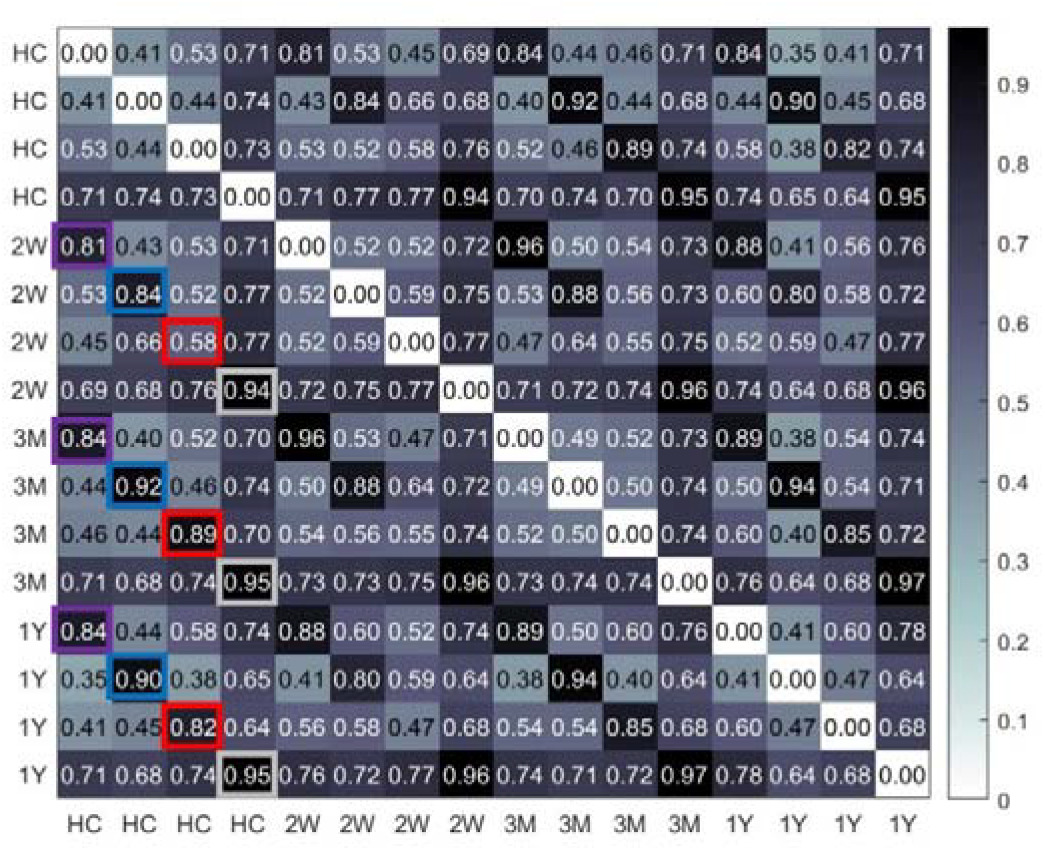
The spatial normalization of cortical language network dynamics. Spatial Pearson correlation coefficients between HCs and patients were calculated for each pair of the four recurring temporal states. Colored rectangles represent state 1 (purple), state 2 (blue), state 3 (red), and state 4 (light gray).

**Figure 6.**
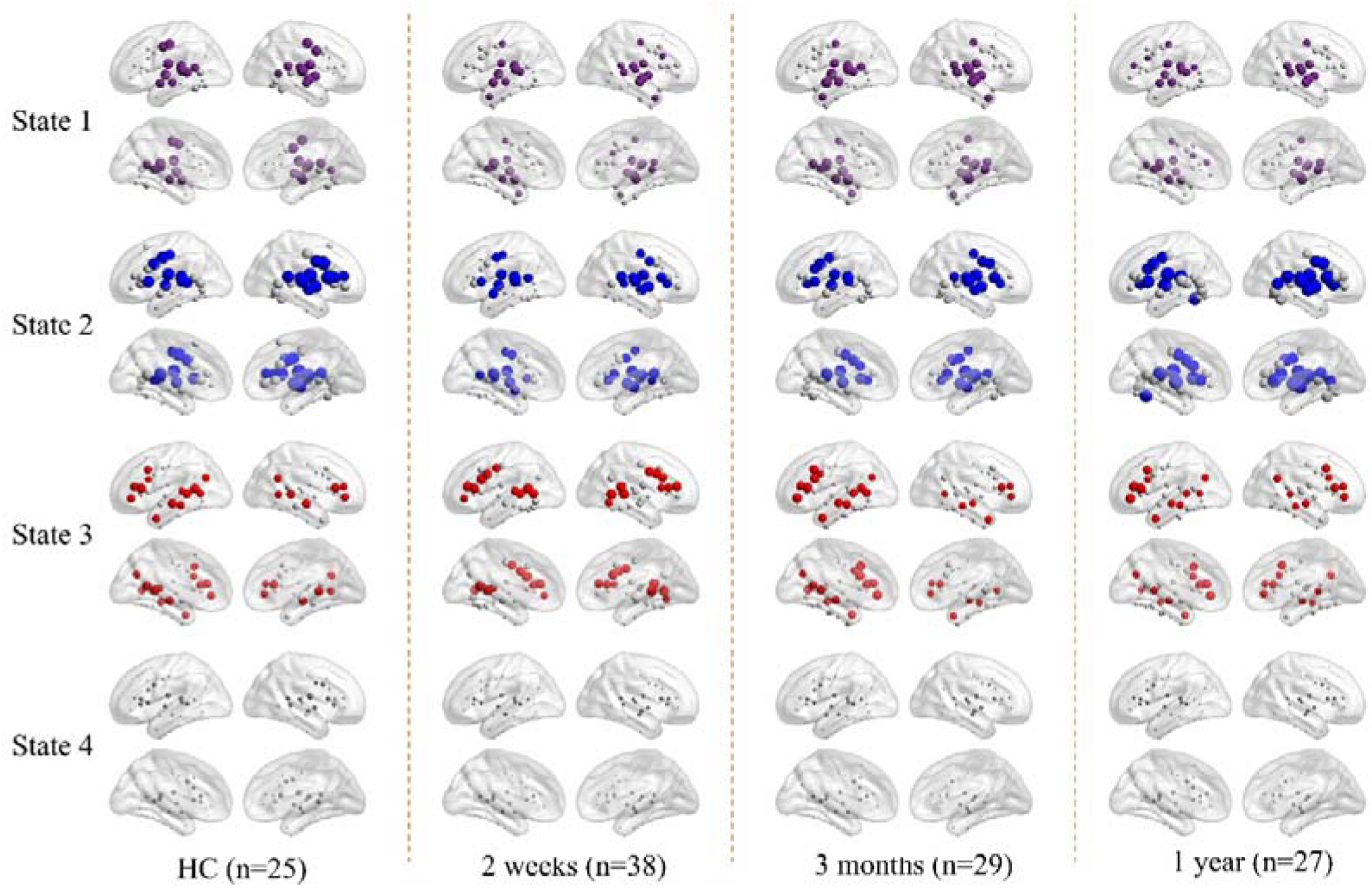
The state-dependent hub distributions. During the acute phase, hubs within the bilateral IFG in state 2 and the anterior temporal cortex in state 3 experienced significant disruption. By 3 months post-stroke, these hubs reappeared, corresponding with observed improvements in language function. The top 20 nodes with the highest nodal strength in each state were colored and designated as hubs.

**Figure 7.**
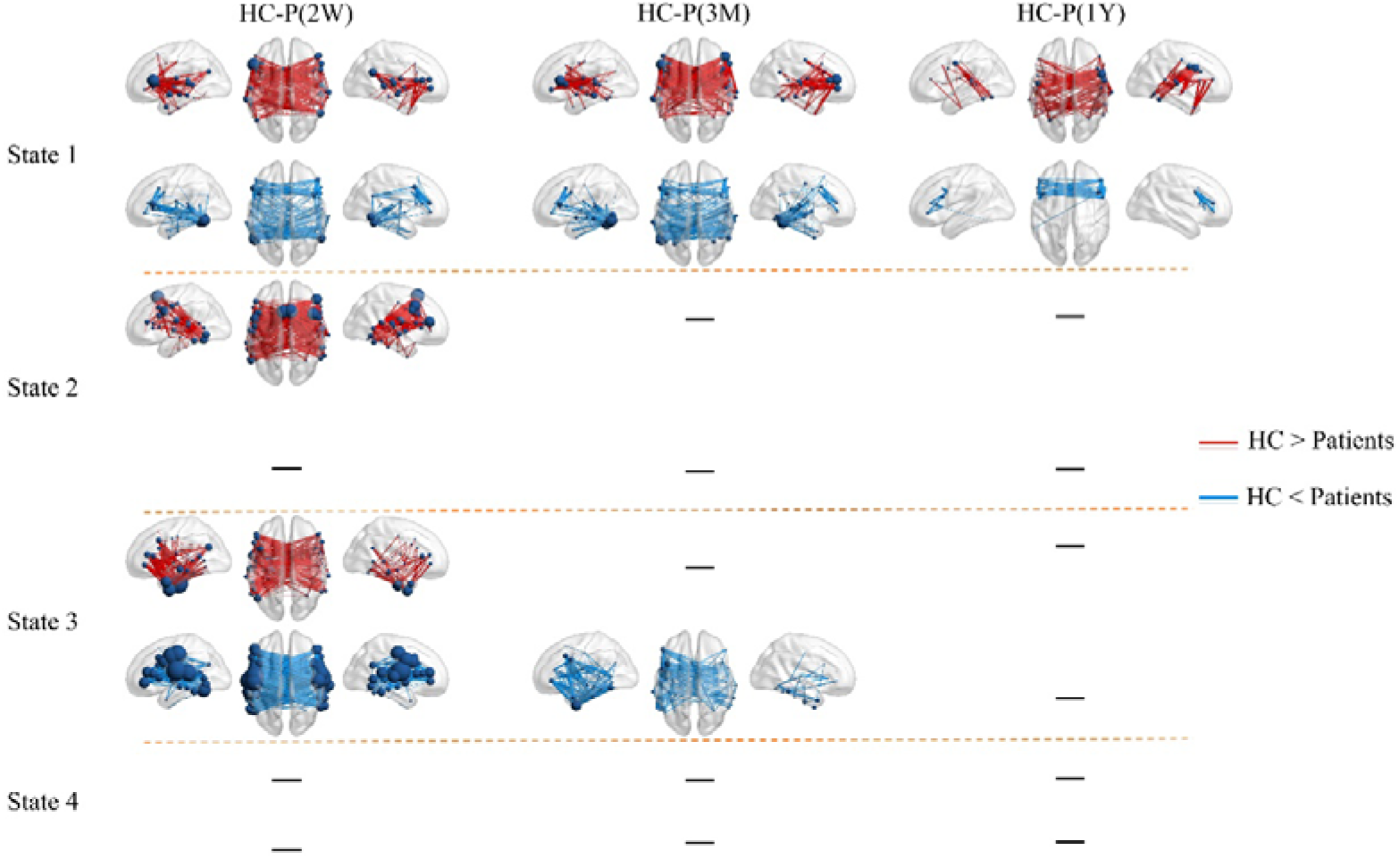
State-dependent network disruptions during the acute phase and normalization during language recovery. Edge *P* <.05, component *P* <.05 with NBS correction.

**Figure 8.**
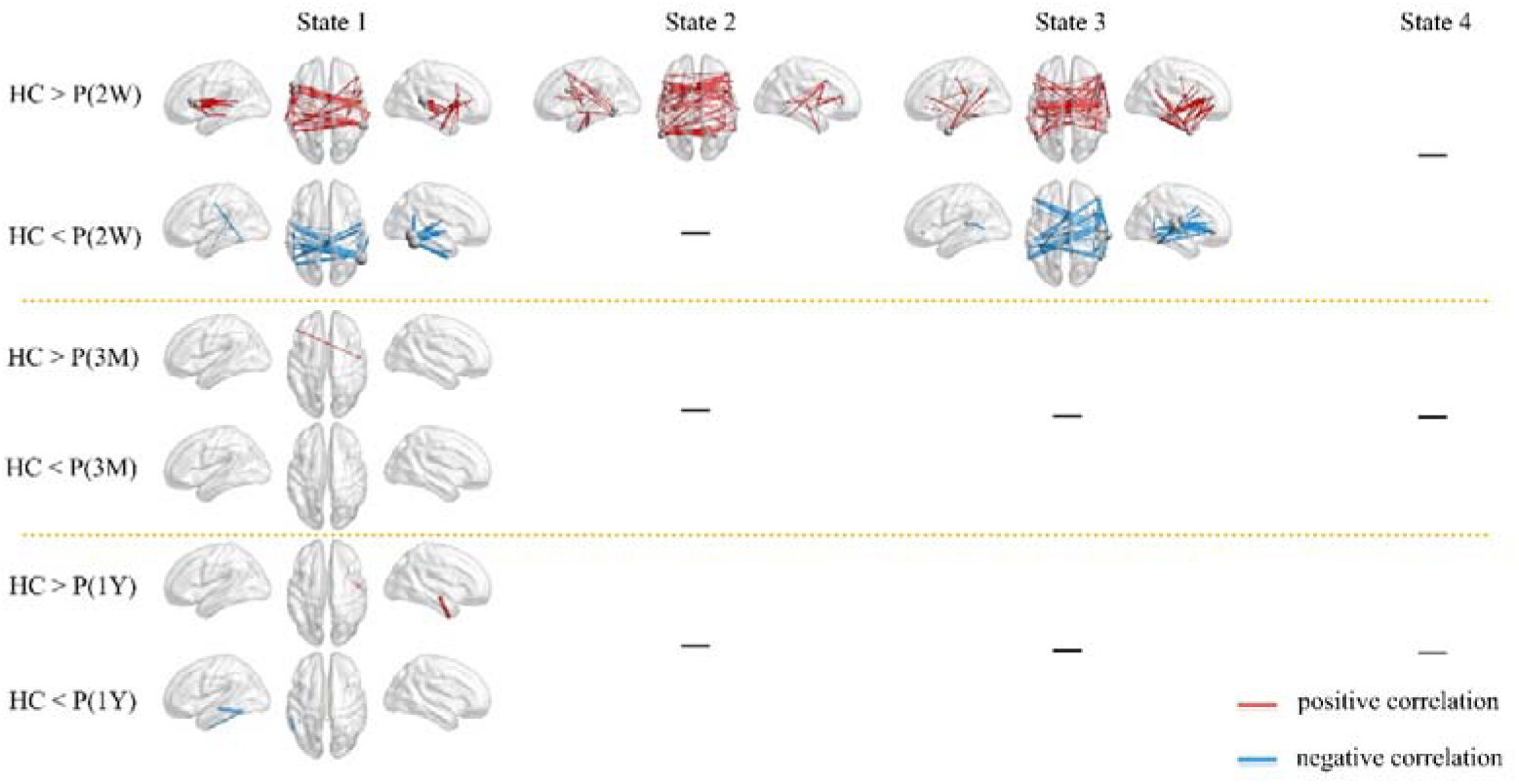
The partial correlation between network disruptions and PCA-based language scores in the acute phase. Edge *P*<.05, component *P*<.05 with NBS correction.

State-specific increases in nodal and global properties were observed during the acute phase (Figures 9 and 10). State 3 exhibited the most significant rises in nodal DC, gE, and lE in the posterior temporal cortex and ventral parts of PrG and PoG. Nodal properties returned to normal levels three months and one year after stroke. State 3 also showed the most significant increases in total strength and both global and local network efficiencies.

**Figure 9.**
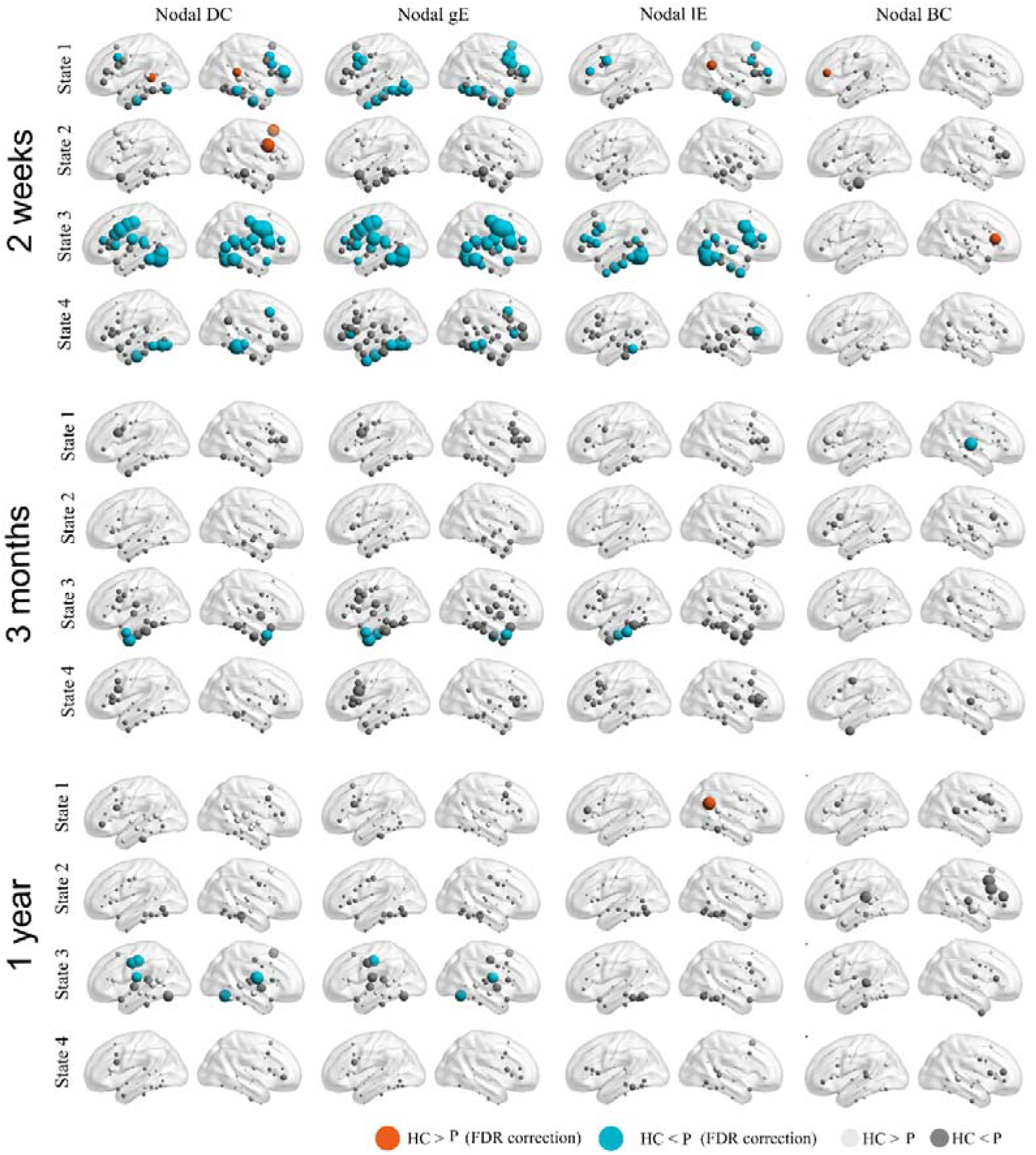
Changes of nodal properties in the acute phase and normalization during language recovery. The results were corrected in each state using FDR with a *P* value of 0.05.

**Figure 10.**
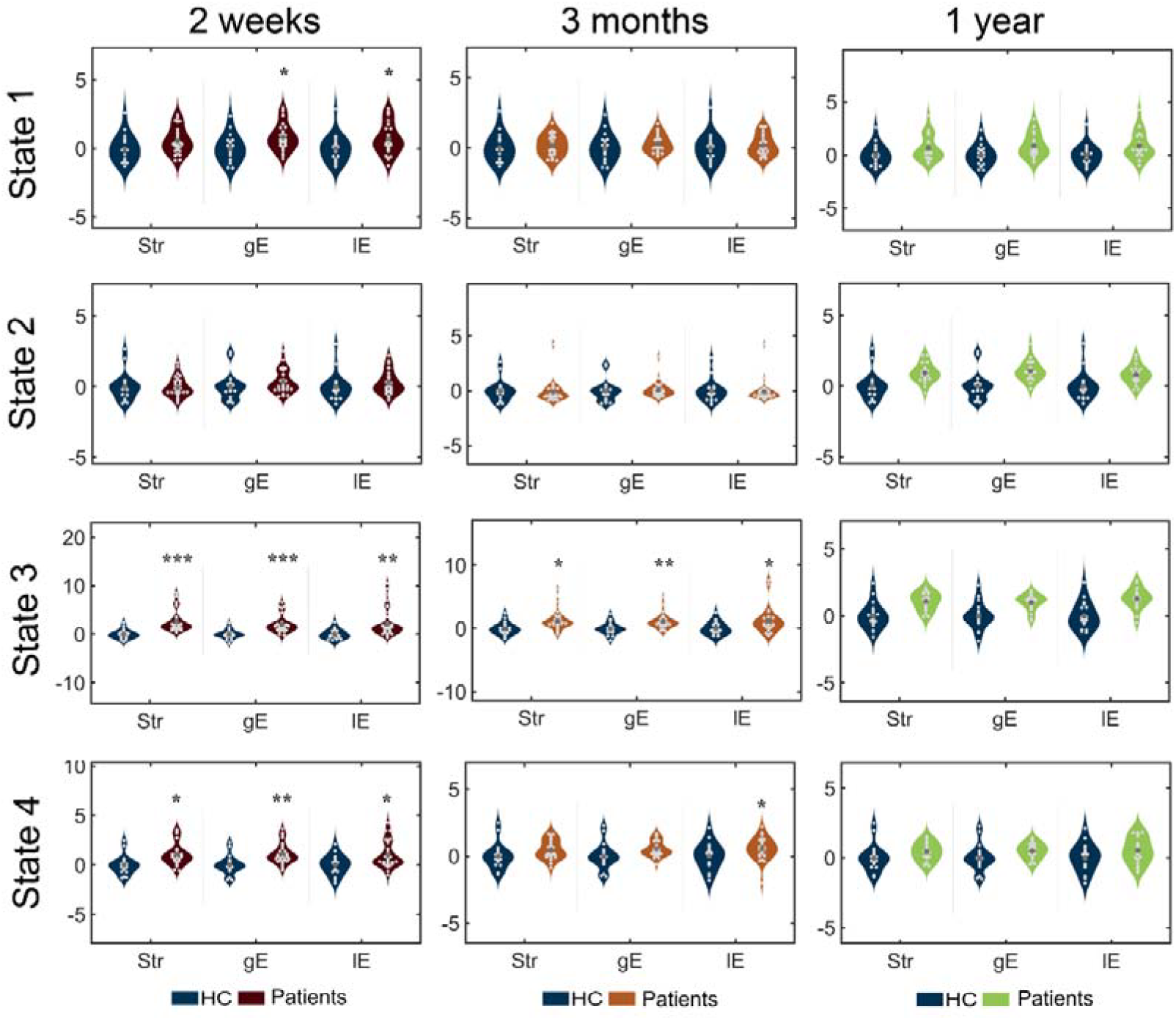
Changes of global properties in the acute phase and normalization during language recovery. ^*^ *P* < 0.05; ^**^, *P* < 0.01; ^***^ *P* < 0.001.

State-specific increases in global network properties were also observed (Figure 10). State 3 showed the most significant increases in global strength, gE, and lE. These global properties normalized toward HCs during language recovery.

Partial analysis revealed no significant link between nodal properties and language deficits or between global properties and language deficits.

## Discussion

The language network consists of a series of cortical regions and their underlying white matter tracks. Disconnection of white matter disrupts the functional interactions between cortical areas, leading to language impairments. In this study, we screened post-stroke patients with pure white matter lesions and examined how white matter disconnection affected the dynamics of the cortical language network. Longitudinal behavioral and rs-fMRI data were collected from 38 patients after subcortical stroke. Unlike previous studies, we utilized the “dynamic meta-networking framework of language” to assess the impact of subcortical stroke on domain-segregation within language sub-networks. The main findings are as follows: First, during the acute phase, the domain-segregation pattern of cortical language dynamics was severely disrupted, impacting state 1 (responsible for phonological perception and representation), state 2 (responsible for speech production), and state 3 (responsible for semantic representation), which correlated with patients’ speech and perception impairments. Second, the language network in the acute phase showed both hypo- and hyper-connectivity, with hypo-connectivity positively associated and hyper-connectivity negatively associated with language deficits. Both disruption patterns were mainly interhemispheric, indicating that subcortical stroke impairs interhemispheric inhibitory interactions. Finally, the cortical language network demonstrated significant neuroplastic potential. By three months, patients exhibited rapid recovery of language functions. As language recovery progressed, the connectivity patterns of the language network quickly returned to normal levels similar to those of healthy controls, and domain-segregation dynamics were re-established. These findings enhance our understanding of how white matter damage impacts the cortical language network and emphasize its considerable neuroplastic potential.

One of the most fundamental characteristics of brain networks is that, within a limited and fixed structural neurobiology, functional networks are divided into functionally specific sub-networks (Khambhati, Sizemore, Betzel, & Bassett, 2018). Complex cognition results from the dynamic interactions among these sub-networks (Herbet & Duffau, 2020; Kulkarni & Bassett, 2025). In the context of language networks, classic and dual-pathway models split complex language processes into functionally specific pathways based on organized components (Hickok, 2022). In this study, we replicated findings from Yuan et al. (2023), showing that both healthy controls and stroke patients in recovery phases (at three months or one year post-stroke) display a domain-segregation dynamic reorganization pattern in the cortical language network. Contrary to classic models and the dual-pathway model, we identify three distinct sub-networks. Specifically, the dorsal pathway in the dual-stream model is divided into the phonological perception and representation pathway (state 1) and the speech production pathway (state 2). The phonological perception and representation pathway is involved not only in expressive language but also in receptive language (Binder, 2017; Hickok, 2022). The independence of State 2 allows it to flexibly connect with States 1 and 3. For example, in listening comprehension tasks, States 1 and 3 are involved sequentially. For speech production tasks, States 3, 1, and 2 are engaged sequentially. In repetition tasks, States 1 and 2 are involved in sequence (Hickok, 2022).

During the acute phase, we observed that the most severely affected were the first three states. In the study by Yuan et al. (2023), they found that the white matter tracts connected to hubs in State 1 are mainly the MdLF, in State 2 are mainly the AF, and in State 3 are mainly the ILF, AF, and IFOF. The analysis of the correlation between white matter disconnection and language impairment showed significant negative correlations with the IFOF, MdLF, and AF (Table 3). This explains why the first three states are heavily disrupted. Additionally, we found that the strongest link between white matter disconnection and language impairment was with the corpus callosum (*r* = -0.735), which also clarifies why the most affected connections in the first three states are those between the hemispheres. These results align with the findings by Langen et al. (2017), which showed that white matter tracts influence the dynamic reorganization of the cortical language network and that damage to these tracts reduces tract-specific functional connectivity.

We also observed that, as language recovers, the language network disrupted during the acute phase quickly reorganizes, returning to a pattern similar to that of the healthy control group, known as the domain-segregation pattern. This suggests that in adult patients with brain injury, the language network demonstrates strong neural plasticity (Herbet, Maheu, Costi, Lafargue, & Duffau, 2016; B. Yuan, Xie, Gong, et al., 2023; Zhang et al., 2021). Neural plasticity is a key neural mechanism underlying functional recovery after brain injury (Stefaniak, Halai, & Lambon Ralph, 2020). Although damage to white matter tracts is irreversible, it mainly affects direct connections (Langen et al., 2017). The functional networks connecting brain regions include not only direct connections but also indirect connection patterns. Evidence strongly suggests that network reorganization does not necessarily rely on the structural integrity of white matter (Li et al., 2019).

This study has several limitations. First, although 38 samples were included, there is considerable variability in the lesions, which involve both hemispheres and several white matter tracts (see Figure 1). Interestingly, some white matter tracts that were not previously thought to be involved in language processing were also found to have significant correlations with language impairment (see Table 3). Whether these white matter tracts directly participate in language processing or influence language abilities by affecting other cognitive functions (such as attention and memory) remains to be further explored. Additionally, subcortical damage includes white matter and parts of the thalamus and basal ganglia. Many studies have demonstrated that subregions of the thalamus and specific nuclei in the basal ganglia are directly involved in language processing (Turker, Kuhnke, Eickhoff, Caspers, & Hartwigsen, 2023), and there are complex functional interactions between these subcortical regions and cortical language areas (Favaretto et al., 2022). Thus, damage observed in the cortical language network cannot exclude these factors.

In summary, based on the dynamic meta-networking framework of language, this study demonstrates how white matter damage affects cortical network reorganization. The impact of white matter disconnection on the cortical language network is temporary. The adult brain shows strong neural plasticity, and as language recovers, these state-specific network disruptions are reestablished during the recovery process.

## Supporting information

SubcorticalStroke_whitematter_supplementary

## CRediT authorship contribution statement

Tao Zhong (Methodology, Formal analysis, Visualization, Writing—original draft), Qingwen Chen (Methodology, Formal analysis, Visualization, Writing—original draft), Xiaolin Guo (Methodology, Formal analysis, Visualization, Writing—original draft), Junjie Yang (Writing— review & editing, Writing—original draft, Visualization), Xiaowei Gao (Writing—review & editing, Writing—original draft, Visualization), Zhe Hu (Writing—review & editing, Writing— original draft, Visualization), Jian Liu (Writing—review & editing, Writing—original draft, Visualization), Junjing Li (Writing—review & editing, Writing—original draft, Visualization), Jiaxuan Liu (Writing—review & editing, Writing—original draft, Visualization), Yaling Wang (Writing—review & editing, Writing—original draft, Visualization), Zhiheng Qu (Writing—review & editing, Writing—original draft, Visualization), Wanchun Li (Writing— review & editing, Writing—original draft, Visualization), Zhongqi Li (Writing—review & editing, Writing—original draft, Visualization), Wanjing Li (Writing—review & editing, Writing—original draft, Visualization), Yien Huang (Writing—review & editing, Writing—original draft, Visualization), Jiali Chen (Writing—review & editing, Writing—original draft, Visualization), Hao Wen (Writing—review & editing, Writing—original draft, Visualization), Binke Yuan (Conceptualization, Methodology, Formal analysis, Investigation, Writing—review & editing, Writing—original draft, Visualization), Han Gao (Conceptualization, Methodology, Formal analysis, Investigation, Writing—review & editing, Writing— original draft, Visualization)

## Data and Code availability

The behavioral and neuroimaging data are publicly available at https://cnda.wustl.edu/data/projects/CCIR_00299. The DCC toolbox was available at https://github.com/yuanbinke/Naturalistic-Dynamic-Network-Toolbox (Yang et al., 2025). Other software used in the study was based on DPABI (http://rfmri.org/dpabi), SPM (http://www.fl.ion.ucl.ac.uk/spm), GRETNA (https://www.nitrc.org/projects/gretna/) and BrainNetViewer (https://www.nitrc.org/projects/bnv/).

## Conflict of Interest

No competing financial interests exist.

## Funding

The study is supported by the National Social Science Foundation of China (No. 20&ZD296), the Key-Area Research and Development Program of Guangdong Province (No. 2019B030335001), the National Natural Science Foundation of China (No.32100889), Research Center for Brain Cognition and Human Development, Guangdong, China (No. 2024B0303390003), Medical Scientific Research Foundation of Guangdong Province, China (No. B2025292), and Plan on enhancing scientific research in GMU, Guangdong, China (GMUCR2025-02028).

## Notes

### Competing Interest Statement

The authors have declared no competing interest.

## References

Abbasi, O., Steingraber, N., Chalas, N., Kluger, D. S., & Gross, J. (2023). Spatiotemporal dynamics characterise spectral connectivity profiles of continuous speaking and listening. PLoS Biol, 21(7), e3002178. Retrieved from https://www.ncbi.nlm.nih.gov/pubmed/37478152. doi:10.1371/journal.pbio.3002178

Allen, E. A., Damaraju, E., Plis, S. M., Erhardt, E. B., Eichele, T., & Calhoun, V. D. (2014). Tracking whole-brain connectivity dynamics in the resting state. Cereb Cortex, 24(3), 663–676. Retrieved from https://www.ncbi.nlm.nih.gov/pubmed/23146964. doi:10.1093/cercor/bhs352

Binder, J. R. (2017). Current Controversies on Wernicke’s Area and its Role in Language. Curr Neurol Neurosci Rep, 17(8), 58. Retrieved from https://www.ncbi.nlm.nih.gov/pubmed/28656532. doi:10.1007/s11910-017-0764-8

Chai, L. R., Mattar, M. G., Blank, I. A., Fedorenko, E., & Bassett, D. S. (2016). Functional Network Dynamics of the Language System. Cereb Cortex, 26(11), 4148–4159. Retrieved from https://www.ncbi.nlm.nih.gov/pubmed/27550868. doi:10.1093/cercor/bhw238

Corbetta, M., Ramsey, L., Callejas, A., Baldassarre, A., Hacker, C. D., Siegel, J. S., Shulman, G. L. (2015). Common behavioral clusters and subcortical anatomy in stroke. Neuron, 85(5), 927–941. Retrieved from https://www.ncbi.nlm.nih.gov/pubmed/25741721. doi:10.1016/j.neuron.2015.02.027

Damaraju, E., Allen, E. A., Belger, A., Ford, J. M., McEwen, S., Mathalon, D. H., Calhoun, V. D. (2014). Dynamic functional connectivity analysis reveals transient states of dysconnectivity in schizophrenia. Neuroimage Clin, 5, 298–308. Retrieved from https://www.ncbi.nlm.nih.gov/pubmed/25161896. doi:10.1016/j.nicl.2014.07.003

Dockes, J., Poldrack, R. A., Primet, R., Gozukan, H., Yarkoni, T., Suchanek, F., Varoquaux, G. (2020). NeuroQuery, comprehensive meta-analysis of human brain mapping. Elife, 9. Retrieved from https://www.ncbi.nlm.nih.gov/pubmed/32129761. doi:10.7554/eLife.53385

Duffau, H., Moritz-Gasser, S., & Mandonnet, E. (2014). A re-examination of neural basis of language processing: proposal of a dynamic hodotopical model from data provided by brain stimulation mapping during picture naming. Brain Lang, 131, 1–10. Retrieved from https://www.ncbi.nlm.nih.gov/pubmed/23866901. doi:10.1016/j.bandl.2013.05.011

Fan, L., Li, H., Zhuo, J., Zhang, Y., Wang, J., Chen, L., Jiang, T. (2016). The Human Brainnetome Atlas: A New Brain Atlas Based on Connectional Architecture. Cereb Cortex, 26(8), 3508–3526. Retrieved from https://www.ncbi.nlm.nih.gov/pubmed/27230218. doi:10.1093/cercor/bhw157

Favaretto, C., Allegra, M., Deco, G., Metcalf, N. V., Griffis, J. C., Shulman, G. L., Corbetta, M. (2022). Subcortical-cortical dynamical states of the human brain and their breakdown in stroke. Nat Commun, 13(1), 5069. Retrieved from https://www.ncbi.nlm.nih.gov/pubmed/36038566. doi:10.1038/s41467-022-32304-1

Fedorenko, E., & Thompson-Schill, S. L. (2014). Reworking the language network. Trends Cogn Sci, 18(3), 120–126. Retrieved from https://www.ncbi.nlm.nih.gov/pubmed/24440115. doi:10.1016/j.tics.2013.12.006

Griffis, J. C., Metcalf, N. V., Corbetta, M., & Shulman, G. L. (2019). Structural Disconnections Explain Brain Network Dysfunction after Stroke. Cell Rep, 28(10), 2527–2540e2529. Retrieved from https://www.ncbi.nlm.nih.gov/pubmed/31484066. doi:10.1016/j.celrep.2019.07.100

Griffis, J. C., Metcalf, N. V., Corbetta, M., & Shulman, G. L. (2021). Lesion Quantification Toolkit: A MATLAB software tool for estimating grey matter damage and white matter disconnections in patients with focal brain lesions. Neuroimage Clin, 30, 102639. Retrieved from https://www.ncbi.nlm.nih.gov/pubmed/33813262. doi:10.1016/j.nicl.2021.102639

He, X., Bassett, D. S., Chaitanya, G., Sperling, M. R., Kozlowski, L., & Tracy, J. I. (2018). Disrupted dynamic network reconfiguration of the language system in temporal lobe epilepsy. Brain, 141(5), 1375–1389. Retrieved from https://www.ncbi.nlm.nih.gov/pubmed/29554279. doi:10.1093/brain/awy042

Herbet, G., & Duffau, H. (2020). Revisiting the Functional Anatomy of the Human Brain: Toward a Meta-Networking Theory of Cerebral Functions. Physiol Rev, 100(3), 1181–1228. Retrieved from https://www.ncbi.nlm.nih.gov/pubmed/32078778. doi:10.1152/physrev.00033.2019

Herbet, G., Maheu, M., Costi, E., Lafargue, G., & Duffau, H. (2016). Mapping neuroplastic potential in brain-damaged patients. Brain, 139(Pt 3), 829–844. Retrieved from https://www.ncbi.nlm.nih.gov/pubmed/26912646. doi:10.1093/brain/awv394

Hickok, G. (2022). The dual stream model of speech and language processing. Handb Clin Neurol, 185, 57–69. Retrieved from https://www.ncbi.nlm.nih.gov/pubmed/35078610. doi:10.1016/B978-0-12-823384-9.00003-7

Honey, C. J., Sporns, O., Cammoun, L., Gigandet, X., Thiran, J. P., Meuli, R., & Hagmann, P. (2009). Predicting human resting-state functional connectivity from structural connectivity. Proc Natl Acad Sci U S A, 106(6), 2035–2040. Retrieved from https://www.ncbi.nlm.nih.gov/pubmed/19188601. doi:10.1073/pnas.0811168106

Khambhati, A. N., Sizemore, A. E., Betzel, R. F., & Bassett, D. S. (2018). Modeling and interpreting mesoscale network dynamics. Neuroimage, 180(Pt B), 337–349. Retrieved from https://www.ncbi.nlm.nih.gov/pubmed/28645844. doi:10.1016/j.neuroimage.2017.06.029

Kulkarni, S., & Bassett, D. S. (2025). Toward Principles of Brain Network Organization and Function. Annu Rev Biophys, 54(1), 353–378. Retrieved from https://www.ncbi.nlm.nih.gov/pubmed/39952667. doi:10.1146/annurev-biophys-030722-110624

Langen, C. D., Zonneveld, H. I., White, T., Huizinga, W., Cremers, L. G. M., de Groot, M., Vernooij, M. W. (2017). White matter lesions relate to tract-specific reductions in functional connectivity. Neurobiol Aging, 51, 97–103. Retrieved from https://www.ncbi.nlm.nih.gov/pubmed/28063366. doi:10.1016/j.neurobiolaging.2016.12.004

Lebo, M. J., & Box-Steffensmeier, J. M. (2008). Dynamic conditional correlations in political science. American Journal of Political Science. doi:52(3), 688–704

Li, Q., Dong, J. W., Del Ferraro, G., Petrovich Brennan, N., Peck, K. K., Tabar, V., Holodny, A. I. (2019). Functional Translocation of Broca’s Area in a Low-Grade Left Frontal Glioma: Graph Theory Reveals the Novel, Adaptive Network Connectivity. Front Neurol, 10, 702. Retrieved from https://www.ncbi.nlm.nih.gov/pubmed/31333562. doi:10.3389/fneur.2019.00702

Liljestrom, M., Kujala, J., Stevenson, C., & Salmelin, R. (2015). Dynamic reconfiguration of the language network preceding onset of speech in picture naming. Hum Brain Mapp, 36(3), 1202–1216. Retrieved from https://www.ncbi.nlm.nih.gov/pubmed/25413681. doi:10.1002/hbm.22697

Lindquist, M. A., Xu, Y., Nebel, M. B., & Caffo, B. S. (2014). Evaluating dynamic bivariate correlations in resting-state fMRI: a comparison study and a new approach. Neuroimage, 101, 531–546. Retrieved from https://www.ncbi.nlm.nih.gov/pubmed/24993894. doi:10.1016/j.neuroimage.2014.06.052

Lipkin, B., Tuckute, G., Affourtit, J., Small, H., Mineroff, Z., Kean, H., Siegelman, M. (2022). Probabilistic atlas for the language network based on precision fMRI data from> 800 individuals. Sci Data, 9(1), 529.

Liu, X., Tu, L., Chen, X., Zhong, M., Niu, M., Zhao, L., Huang, R. (2020). Dynamic Language Network in Early and Late Cantonese-Mandarin Bilinguals. Front Psychol, 11, 1189. Retrieved from https://www.ncbi.nlm.nih.gov/pubmed/32625136. doi:10.3389/fpsyg.2020.01189

Lu, J., Zhao, Z., Zhang, J., Wu, B., Zhu, Y., Chang, E. F., Berger, M. S. (2021). Functional maps of direct electrical stimulation-induced speech arrest and anomia: a multicentre retrospective study. Brain, 144(8), 2541–2553. Retrieved from https://www.ncbi.nlm.nih.gov/pubmed/33792674. doi:10.1093/brain/awab125

Marini, A., Galetto, V., Zampieri, E., Vorano, L., Zettin, M., & Carlomagno, S. (2011). Narrative language in traumatic brain injury. Neuropsychologia, 49(10), 2904–2910. Retrieved from https://www.ncbi.nlm.nih.gov/pubmed/21723304. doi:10.1016/j.neuropsychologia.2011.06.017

Messe, A. (2020). Parcellation influence on the connectivity-based structure-function relationship in the human brain. Hum Brain Mapp, 41(5), 1167–1180. Retrieved from https://www.ncbi.nlm.nih.gov/pubmed/31746083. doi:10.1002/hbm.24866

Murphy, K., & Fox, M. D. (2017). Towards a consensus regarding global signal regression for resting state functional connectivity MRI. Neuroimage, 154, 169–173. Retrieved from https://www.ncbi.nlm.nih.gov/pubmed/27888059. doi:10.1016/j.neuroimage.2016.11.052

Nozais, V., Forkel, S. J., Petit, L., Talozzi, L., Corbetta, M., Thiebaut de Schotten, M., & Joliot, M. (2023). Atlasing white matter and grey matter joint contributions to resting-state networks in the human brain. Commun Biol, 6(1), 726. Retrieved from https://www.ncbi.nlm.nih.gov/pubmed/37452124. doi:10.1038/s42003-023-05107-3

Parsons, N., Hughes, M. E., Poudel, G. R., D., J. F., & Caeyenberghs, K. (2020). Structure-Function Relationships in Brain-Injured Patients: A Scoping Review.

Price, C. J. (2012). A review and synthesis of the first 20 years of PET and fMRI studies of heard speech, spoken language and reading. Neuroimage, 62(2), 816–847. Retrieved from https://www.ncbi.nlm.nih.gov/pubmed/22584224. doi:10.1016/j.neuroimage.2012.04.062

Rubinov, M., & Sporns, O. (2010). Complex network measures of brain connectivity: uses and interpretations. Neuroimage, 52(3), 1059–1069. Retrieved from https://www.ncbi.nlm.nih.gov/pubmed/19819337. doi:10.1016/j.neuroimage.2009.10.003

Salvalaggio, A., De Filippo De Grazia, M., Zorzi, M., Thiebaut de Schotten, M., & Corbetta, M. (2020). Post-stroke deficit prediction from lesion and indirect structural and functional disconnection. Brain, 143(7), 2173–2188. Retrieved from https://www.ncbi.nlm.nih.gov/pubmed/32572442. doi:10.1093/brain/awaa156

Saur, D., Kreher, B. W., Schnell, S., Kummerer, D., Kellmeyer, P., Vry, M. S., Weiller, C. (2008). Ventral and dorsal pathways for language. Proc Natl Acad Sci U S A, 105(46), 18035–18040. Retrieved from https://www.ncbi.nlm.nih.gov/pubmed/19004769. doi:10.1073/pnas.0805234105

Stefaniak, J. D., Halai, A. D., & Lambon Ralph, M. A. (2020). The neural and neurocomputational bases of recovery from post-stroke aphasia. Nat Rev Neurol, 16(1), 43–55. Retrieved from https://www.ncbi.nlm.nih.gov/pubmed/31772339. doi:10.1038/s41582-019-0282-1

Turker, S., Kuhnke, P., Eickhoff, S. B., Caspers, S., & Hartwigsen, G. (2023). Cortical, subcortical, and cerebellar contributions to language processing: A meta-analytic review of 403 neuroimaging experiments. Psychol Bull, 149(11-12), 699–723. Retrieved from https://www.ncbi.nlm.nih.gov/pubmed/37768610. doi:10.1037/bul0000403

van den Heuvel, M. P., Mandl, R. C., Kahn, R. S., & Hulshoff Pol, H. E. (2009). Functionally linked resting-state networks reflect the underlying structural connectivity architecture of the human brain. Hum Brain Mapp, 30(10), 3127–3141. Retrieved from https://www.ncbi.nlm.nih.gov/pubmed/19235882. doi:10.1002/hbm.20737

Vigneau, M., Beaucousin, V., Herve, P. Y., Duffau, H., Crivello, F., Houde, O., Tzourio-Mazoyer, N. (2006). Meta-analyzing left hemisphere language areas: phonology, semantics, and sentence processing. Neuroimage, 30(4), 1414–1432. Retrieved from https://www.ncbi.nlm.nih.gov/pubmed/16413796. doi:10.1016/j.neuroimage.2005.11.002

Wang, Z., Dai, Z., Gong, G., Zhou, C., & He, Y. (2015). Understanding structural-functional relationships in the human brain: a large-scale network perspective. Neuroscientist, 21(3), 290–305. Retrieved from https://www.ncbi.nlm.nih.gov/pubmed/24962094. doi:10.1177/1073858414537560

Yang, J., Hu, Z., Li, J., Guo, X., Gao, X., Liu, J., Yuan, B. (2025). NaDyNet: A toolbox for dynamic network analysis of naturalistic stimuli. Neuroimage, 311, 121203. Retrieved from https://www.ncbi.nlm.nih.gov/pubmed/40221067. doi:10.1016/j.neuroimage.2025.121203

Yeh, F. C. (2022). Population-based tract-to-region connectome of the human brain and its hierarchical topology. Nat Commun, 13(1), 4933. Retrieved from https://www.ncbi.nlm.nih.gov/pubmed/35995773. doi:10.1038/s41467-022-32595-4

Youssofzadeh, V., & Babajani-Feremi, A. (2019). Mapping critical hubs of receptive and expressive language using MEG: A comparison against fMRI. Neuroimage, 201, 116029. Retrieved from https://www.ncbi.nlm.nih.gov/pubmed/31325641. doi:10.1016/j.neuroimage.2019.116029

Yuan, B., Xie, H., Gong, F., Zhang, N., Xu, Y., Zhang, H., Yan, J. (2023). Dynamic network reorganization underlying neuroplasticity: the deficits-severity-related language network dynamics in patients with left hemispheric gliomas involving language network. Cereb Cortex, 33(13), 8273–8285. Retrieved from https://www.ncbi.nlm.nih.gov/pubmed/37005067. doi:10.1093/cercor/bhad113

Yuan, B., Xie, H., Wang, Z., Xu, Y., Zhang, H., Liu, J., Wu, J. (2023). The domain-separation language network dynamics in resting state support its flexible functional segregation and integration during language and speech processing. Neuroimage, 274, 120132. Retrieved from https://www.ncbi.nlm.nih.gov/pubmed/37105337. doi:10.1016/j.neuroimage.2023.120132

Yuan, B., Zhang, N., Gong, F., Wang, X., Yan, J., Lu, J., & Wu, J. (2022). Longitudinal assessment of network reorganizations and language recovery in postoperative patients with glioma. Brain Commun, 4(2), fcac046.

Zalesky, A., Fornito, A., & Bullmore, E. T. (2010). Network-based statistic: identifying differences in brain networks. Neuroimage, 53(4), 1197–1207. Retrieved from https://www.ncbi.nlm.nih.gov/pubmed/20600983. doi:10.1016/j.neuroimage.2010.06.041

Zhang, N., Yuan, B., Yan, J., Cheng, J., Lu, J., & Wu, J. (2021). Multivariate machine learning-based language mapping in glioma patients based on lesion topography. Brain Imaging Behav, 15(5), 2552–2562. Retrieved from https://www.ncbi.nlm.nih.gov/pubmed/33619646. doi:10.1007/s11682-021-00457-0

