## Supplementary material for "The Disruption and Normalization of Cortical Language Network Dynamics After Post-Stroke White Matter Disconnection": SubcorticalStroke_whitematter_supplementary

**Supplementary Figures**

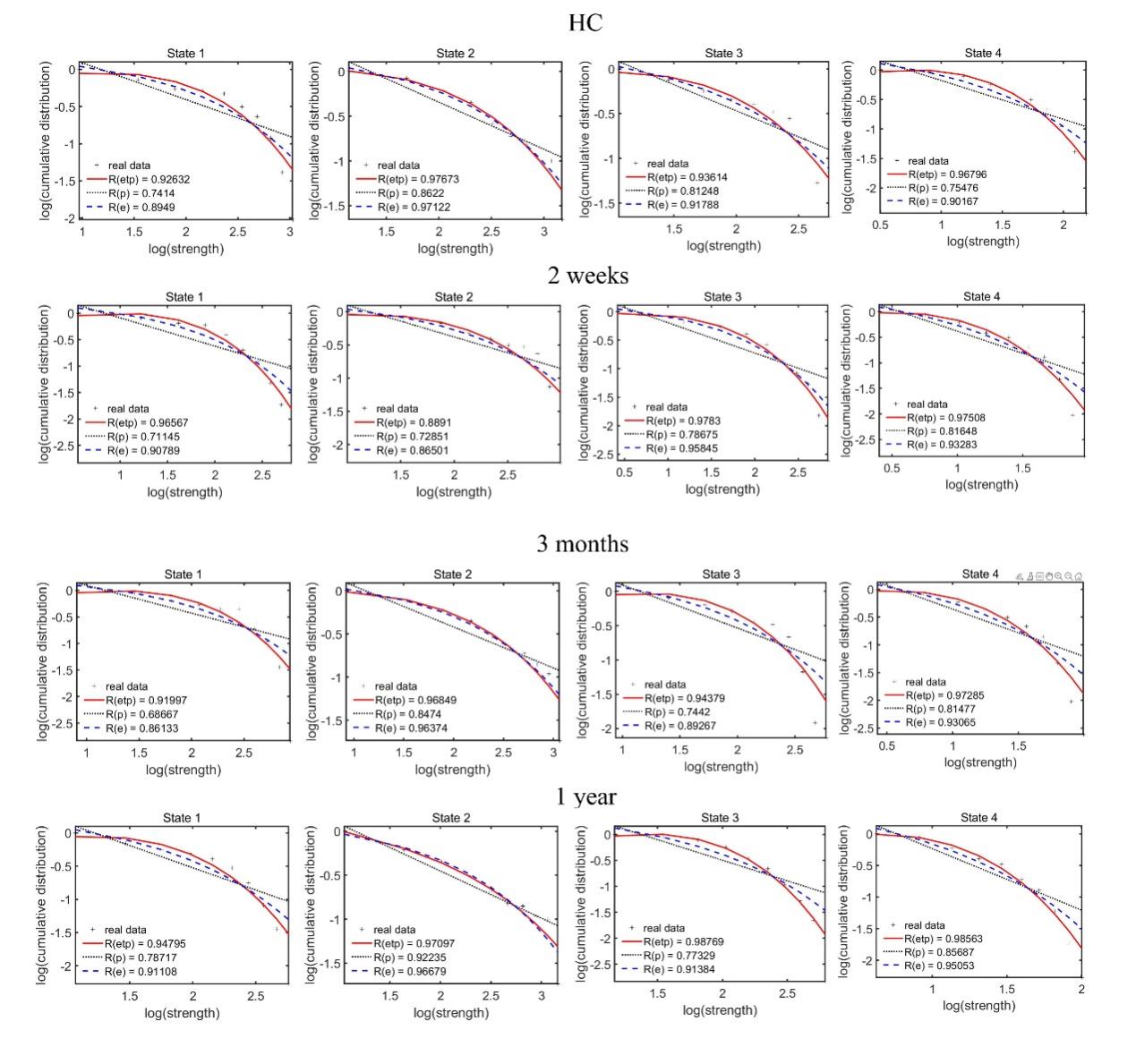

**Supplementary Figure 1.** The log-log plots of the cumulative nodal strength distributions of patients. The plus sign (black) represents observed data, the solid line (red) is the fit of the exponentially truncated power-law, $P \left( x \right) \sim x^{\alpha-1}exp(\frac{x}{x_{c}})$, the dashed line (blue) is an exponential, $P \left( x \right) \sim\exp\left( \frac{x}{x_{c}} \right)$, and the dotted line (black) is a power-law, $P \left( x \right) \sim x^{\alpha-1}.$*R*^2^ was calculated to assess the goodness-of-fit. A larger value indicates a better fitting: *R_etp_*, *R^2^* for the exponentially truncated power-law; *R_e_*, *R^2^* for the exponential; *R_p_*, *R^2^* for the power-law fit. The exponentially truncated power-law is the best fitting for all 4 states, which suggest a long-tailed broad-scale topologies and a large proportion of network connectivity will be concentrated on a subset node (i.e., hubs).

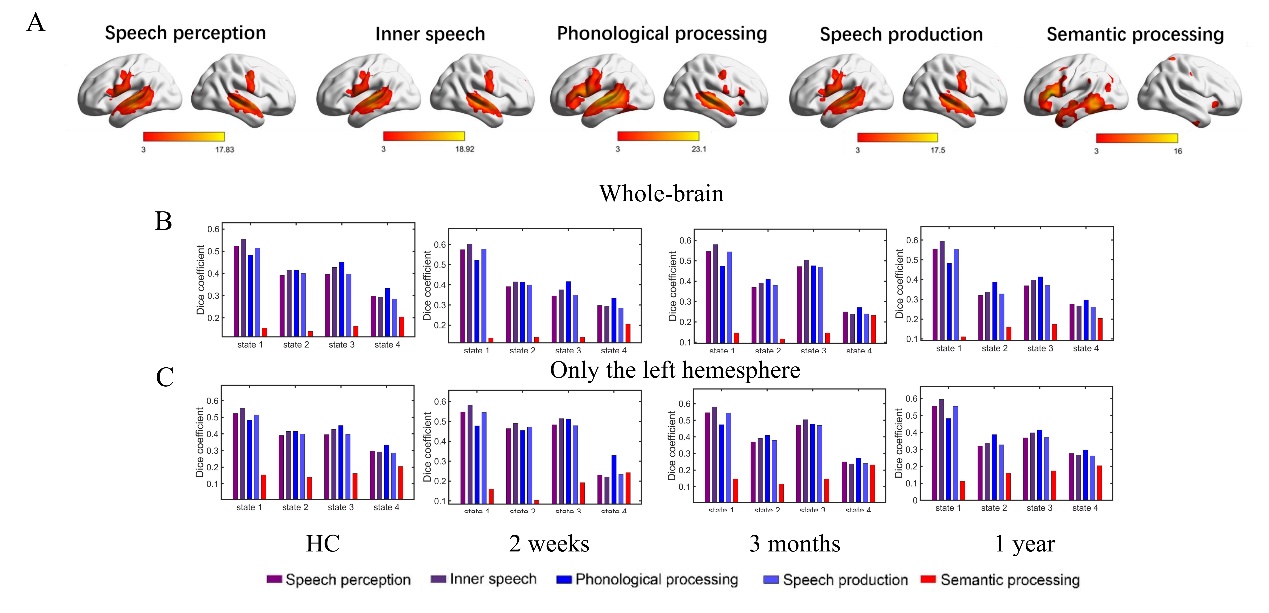

**Supplementary Figure 2.** Functional relevance of hub distributions. A. The meta results of speech perception, inner speech, phonological processing, speech production, and semantic processing were from 'NeuroQuery’ (https://neuroquery.org/) (Dockes et al., 2020). Each map was thresholded at Z = 3 (a typical value used by NeuroQuery) for illustrative purposes and only positive results were shown. B&C: the dice coefficients between binary images of hub nodes and meta results. Considering the left-lateralized activations of meta results, the dice coefficients were calculated at the whole-brain level and in the left hemisphere.
